# Proteomic and Transcriptomic Analyses of the Hippocampus and Cortex in SUDEP and High-Risk SUDEP Cases

**DOI:** 10.1101/2020.07.27.223446

**Authors:** Dominique F. Leitner, James D. Mills, Geoffrey Pires, Arline Faustin, Eleanor Drummond, Evgeny Kanshin, Shruti Nayak, Manor Askenazi, Chloe Verducci, Bei Jun Chen, Michael Janitz, Jasper J. Anink, Johannes C. Baayen, Sander Idema, Erwin A. van Vliet, Sasha Devore, Daniel Friedman, Beate Diehl, Catherine Scott, Roland Thijs, Thomas Wisniewski, Beatrix Ueberheide, Maria Thom, Eleonora Aronica, Orrin Devinsky

**Author notes:** These authors contributed equally to this work (Dominique F. Leitner, James D. Mills, Geoffrey Pires).

## Abstract

Sudden unexpected death in epilepsy (SUDEP) is the leading type of epilepsy-related death. Severely depressed brain activity in these cases may impair respiration, arousal, and protective reflexes, occurring as a prolonged postictal generalized EEG suppression (PGES) and resulting in a high-risk for SUDEP. In autopsy hippocampus and cortex, we observed no proteomic differences between SUDEP and epilepsy cases, contrasting our previously reported robust differences between epilepsy and controls. Transcriptomics in hippocampus and cortex from surgical epilepsy cases segregated by PGES identified 55 differentially expressed genes (37 protein-coding, 15 lncRNAs, three pending) in hippocampus. Overall, the SUDEP proteome and high-risk SUDEP transcriptome largely reflected other epilepsy cases in the brain regions analyzed, consistent with diverse epilepsy syndromes and comorbidities associated with SUDEP. Thus, studies with larger cohorts and different epilepsy syndromes, as well as additional anatomic regions may identify molecular mechanisms of SUDEP.

## Introduction

Epilepsy affects over 65 million people worldwide and markedly increases mortality from direct and indirect effects of seizures and antiseizure therapies (*1*). Sudden unexpected death in epilepsy (SUDEP) affects 1 in 1000 epilepsy patients annually and is the leading cause of epilepsy-related deaths (*2*). SUDEP most often follows a generalized tonic-clonic seizure (GTCS), and excludes trauma, drowning, status epilepticus, or other causes (*3, 4*). Most SUDEP cases are unwitnessed, occur during sleep, and the subject is found prone.

Studies on SUDEP epidemiology, risk factors, mechanisms, and prevention have advanced our understanding, although detailed pathophysiological understanding remains limited (*5–7*). After a GTCS, prolonged (i.e. >50 sec) postictal generalized EEG suppression (PGES) may predispose a patient to SUDEP and may be a SUDEP biomarker, as severe prolonged reduced brain activity impairs arousal, respiration, and other autonomic functions (*8–10*). However, we cannot predict why some low-risk patients suffer SUDEP, high-risk patients survive for decades, and other patients succumb to SUDEP despite recovering from many earlier GTCS. SUDEP cases may harbor pathogenic gene variants in brain and heart ion channels (*11–14*), but a role in SUDEP pathogenesis remains speculative. Animal models of genetic epilepsies and chemo-induced seizures implicate abnormalities in respiration, arousal, and parasympathetic hyperactivity in SUDEP pathogenesis (*2, 15–18*). However, the neuropathology of SUDEP parallels findings in non-SUDEP epilepsy cases (*19, 20*). Potential proteomic and transcriptional molecular signatures associated with SUDEP have not been studied.

Here, we sought to identify molecular signaling networks associated with SUDEP in brain regions implicated in ictogenesis (*21, 22*) using two different approaches. We first evaluated localized protein expression in the microdissected hippocampal CA1-3 region, dentate gyrus, and the superior frontal gyrus from autopsy SUDEP and non-SUDEP epilepsy cases. We also evaluated the transcriptomic differences in hippocampus and temporal cortex of low and high-risk SUDEP (PGES < or ≥ 50 seconds) mesial temporal lobe epilepsy (MTLE) cases from surgical tissue.

## Results

### Proteome of SUDEP and non-SUDEP epilepsy autopsy cases

The differential expression of proteins in autopsy SUDEP (n=12) and non-SUDEP (n=14) epilepsy cases was evaluated using label-free quantitative mass spectrometry (MS) in the microdissected hippocampal CA1-3 region, dentate gyrus, and superior frontal cortex, as these regions have been implicated in ictogenesis and may also be impacted by seizure activity (*21, 22*). Case histories are summarized in Table 1 and Fig. 1A-B. A principal component analysis (PCA) did not distinguish SUDEP and non-SUDEP epilepsy cases in any of the studied brain regions (Fig. 1C-E). The main source of variation in these cases, PCA1, did not show a significant difference when comparing SUDEP and non-SUDEP epilepsy cases in each brain region by an unpaired two-tailed t test, as depicted by a box plot in Fig. 1C-E. Lifetime GTCS burden, associated with an increased SUDEP risk (*1, 2*), was evaluated to determine whether this factor may contribute to protein differences as seen by a separation of groups. From cases with available data (9 SUDEP and 8 non-SUDEP epilepsy cases), 55.6% of SUDEP and 62.5% of non-SUDEP epilepsy cases had > 10 lifetime GTCS, and 22.2% of SUDEP cases and 12.5% of non-SUDEP epilepsy cases had > 100 lifetime GTCS. Lifetime GTCS frequency did not contribute to group differences in the PCA (Fig.1C-E). There was no enrichment in SUDEP or non-SUDEP epilepsy cases with > 10 or > 100 lifetime GTCS by a Fisher’s exact test. Further, in the PCA, there was no relationship of SUDEP status to neuropathology (focal cortical dysplasia (FCD, n = 10), hippocampal dentate gyrus dysgenesis (n = 7), hippocampal sclerosis (n = 3), and gliosis (n = 3)). Of note, microdissected regions did not necessarily contain observed FCD as it may have been present in other brain regions. Similarly, neuropathology was unrelated to SUDEP status (FCD in 50% of SUDEP cases versus 28.6% of non-SUDEP epilepsy cases, Fisher’s exact test, p = 0.4216).

**Table 1.**
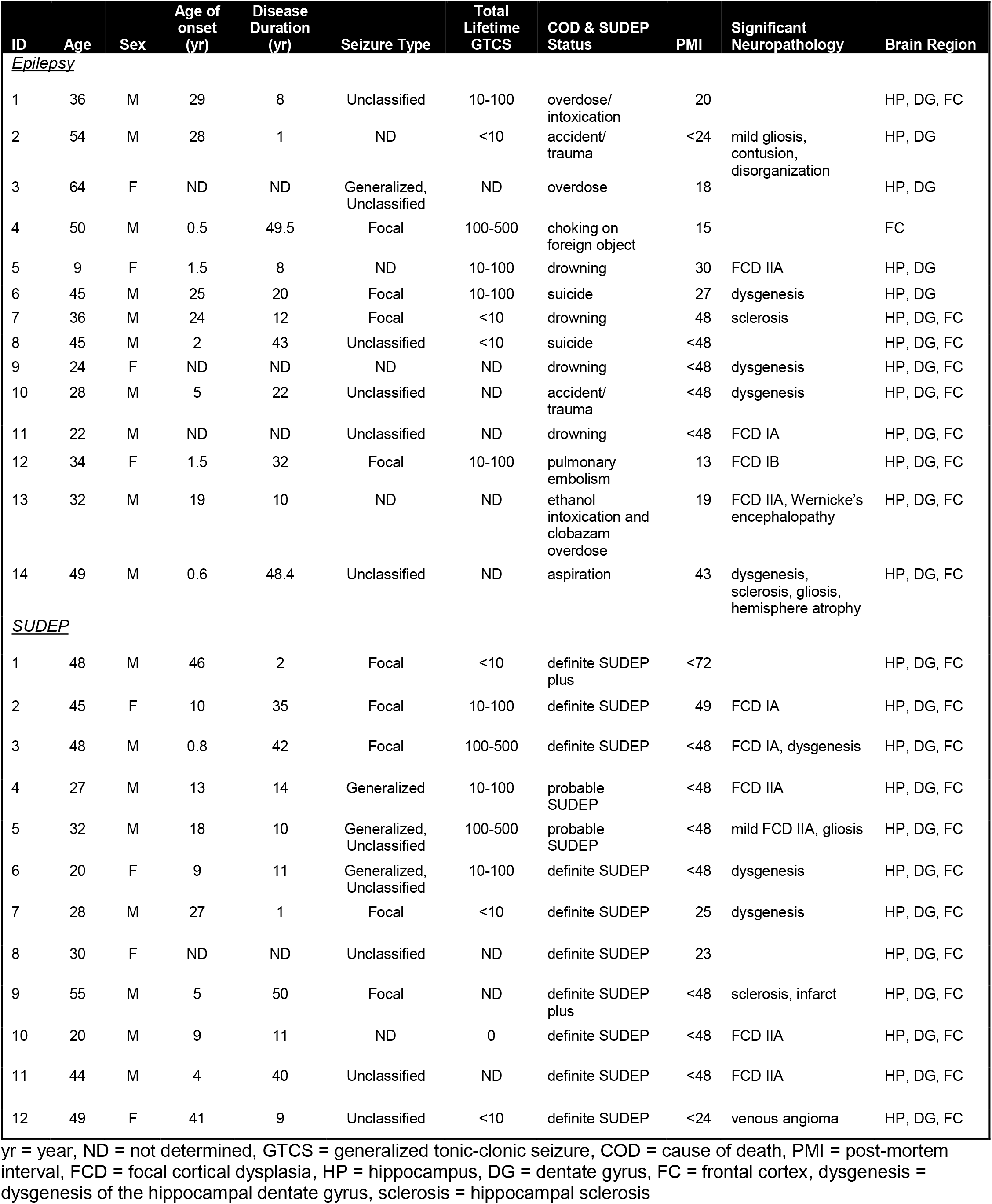
Epilepsy and SUDEP Cases in Proteomics Analyses.

| ID | Age | Sex | Age of onset (yr) | Disease Duration (yr) | Seizure Type | Total Lifetime GTCS | COD & SUDEP Status | PMI | Significant Neuropathology | Brain Region |
| --- | --- | --- | --- | --- | --- | --- | --- | --- | --- | --- |
| <u>*Epilepsy*</u> |  |  |  |  |  |  |  |  |  |  |
| 1 | 36 | M | 29 | 8 | Unclassified | 10-100 | overdose/intoxication | 20 |  | HP, DG, FC |
| 2 | 54 | M | 28 | 1 | ND | <10 | accident/trauma | <24 | mild gliosis, contusion, disorganization | HP, DG |
| 3 | 64 | F | ND | ND | Generalized, Unclassified | ND | overdose | 18 |  | HP, DG |
| 4 | 50 | M | 0.5 | 49.5 | Focal | 100-500 | choking on foreign object | 15 |  | FC |
| 5 | 9 | F | 1.5 | 8 | ND | 10-100 | drowning | 30 | FCD IIA | HP, DG |
| 6 | 45 | M | 25 | 20 | Focal | 10-100 | suicide | 27 | dysgenesis | HP, DG |
| 7 | 36 | M | 24 | 12 | Focal | <10 | drowning | 48 | sclerosis | HP, DG, FC |
| 8 | 45 | M | 2 | 43 | Unclassified | <10 | suicide | <48 |  | HP, DG, FC |
| 9 | 24 | F | ND | ND | ND | ND | drowning | <48 | dysgenesis | HP, DG, FC |
| 10 | 28 | M | 5 | 22 | Unclassified | ND | accident/trauma | <48 | dysgenesis | HP, DG, FC |
| 11 | 22 | M | ND | ND | Unclassified | ND | drowning | <48 | FCD IA | HP, DG, FC |
| 12 | 34 | F | 1.5 | 32 | Focal | 10-100 | pulmonary embolism | 13 | FCD IB | HP, DG, FC |
| 13 | 32 | M | 19 | 10 | ND | ND | ethanol intoxication and clobazam overdose | 19 | FCD IIA, Wernicke's encephalopathy | HP, DG, FC |
| 14 | 49 | M | 0.6 | 48.4 | Unclassified | ND | aspiration | 43 | dysgenesis, sclerosis, gliosis, hemisphere atrophy | HP, DG, FC |
| <u>*SUDEP*</u> |  |  |  |  |  |  |  |  |  |  |
| 1 | 48 | M | 46 | 2 | Focal | <10 | definite SUDEP plus | <72 |  | HP, DG, FC |
| 2 | 45 | F | 10 | 35 | Focal | 10-100 | definite SUDEP | 49 | FCD IA | HP, DG, FC |
| 3 | 48 | M | 0.8 | 42 | Focal | 100-500 | definite SUDEP | <48 | FCD IA, dysgenesis | HP, DG, FC |
| 4 | 27 | M | 13 | 14 | Generalized | 10-100 | probable SUDEP | <48 | FCD IIA | HP, DG, FC |
| 5 | 32 | M | 18 | 10 | Generalized, Unclassified | 100-500 | probable SUDEP | <48 | mild FCD IIA, gliosis | HP, DG, FC |
| 6 | 20 | F | 9 | 11 | Generalized, Unclassified | 10-100 | definite SUDEP | <48 | dysgenesis | HP, DG, FC |
| 7 | 28 | M | 27 | 1 | Focal | <10 | definite SUDEP | 25 | dysgenesis | HP, DG, FC |
| 8 | 30 | F | ND | ND | Unclassified | ND | definite SUDEP | 23 |  | HP, DG, FC |
| 9 | 55 | M | 5 | 50 | Focal | ND | definite SUDEP plus | <48 | sclerosis, infarct | HP, DG, FC |
| 10 | 20 | M | 9 | 11 | ND | 0 | definite SUDEP | <48 | FCD IIA | HP, DG, FC |
| 11 | 44 | M | 4 | 40 | Unclassified | ND | definite SUDEP | <48 | FCD IIA | HP, DG, FC |
| 12 | 49 | F | 41 | 9 | Unclassified | <10 | definite SUDEP | <24 | venous angioma | HP, DG, FC |

yr = year, ND = not determined, GTCS = generalized tonic-clonic seizure, COD = cause of death, PMI = post-mortem interval, FCD = focal cortical dysplasia, HP = hippocampus, DG = dentate gyrus, FC = frontal cortex, dysgenesis = dysgenesis of the hippocampal dentate gyrus, sclerosis = hippocampal sclerosis

**Figure 1.**
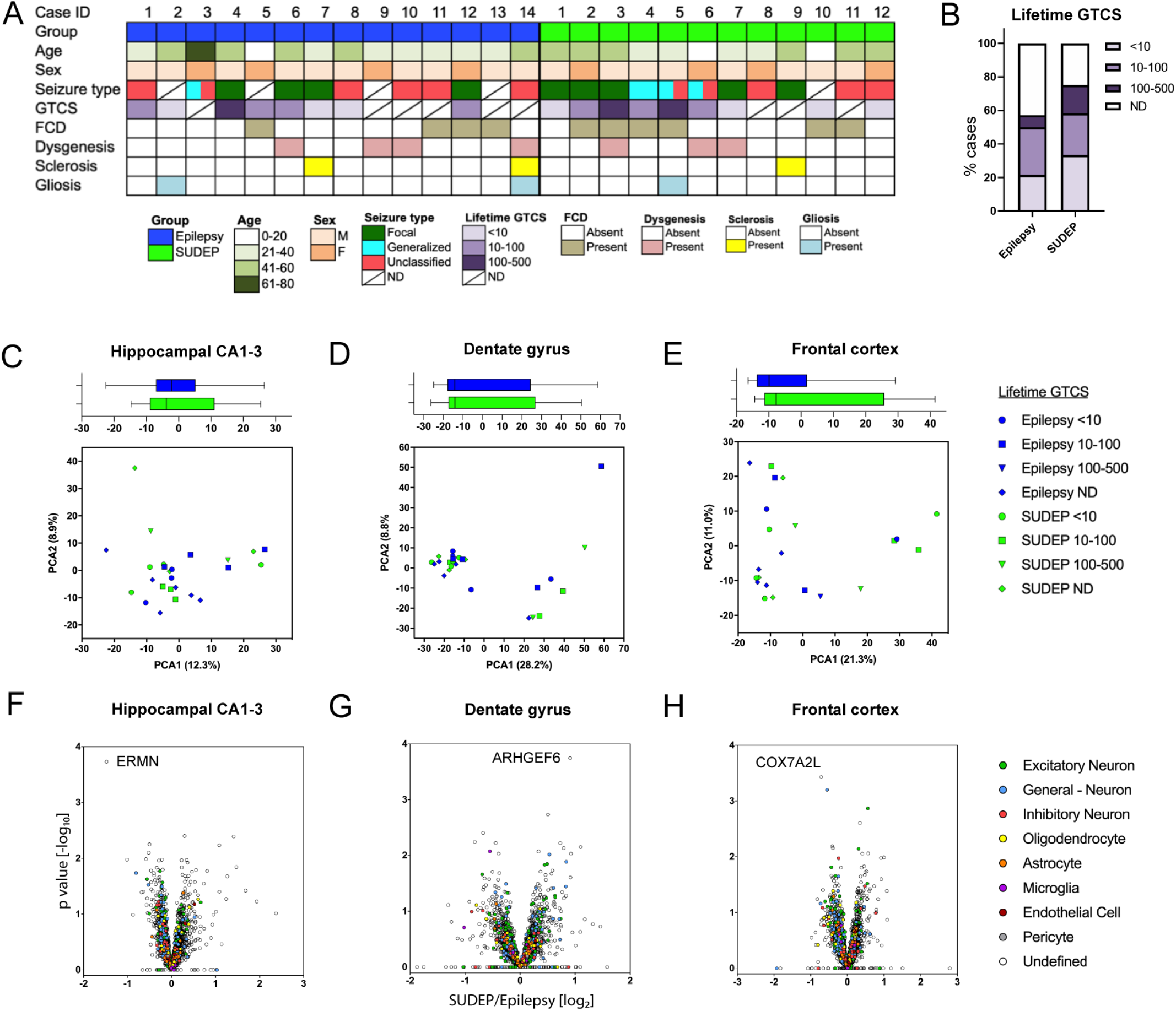
Proteomics analyses in the hippocampus, dentate gyrus, and frontal cortex of SUDEP and non-SUDEP epilepsy cases. **A)** Case history summary for autopsy SUDEP and non-SUDEP epilepsy cases. **B)** A summary of lifetime GTCS history burden for the cases in this study. **C-E)** A principal component analysis (PCA) of the proteomics analyses shows the indicated variation in each brain region of SUDEP cases (n=12) and non-SUDEP epilepsy cases (n=14). There is no separation by SUDEP status or lifetime GTCS history burden. An unpaired two-tailed t test of PCA1 between the SUDEP and non-SUDEP epilepsy groups in each brain region was not significant, as depicted by a box plot with bars indicating minimum and maximum values. ND = not determined. **F-H)** Volcano plots indicate that there are no significantly different proteins in the hippocampal CA1-3 region, dentate gyrus, or frontal cortex of SUDEP and non-SUDEP epilepsy cases as determined by a student’s two tailed t-test with permutation correction at a 5% FDR. The top proteins with the lowest p values in each brain region are noted. Cell type specific protein annotation is included, with the most predominant listed in decreasing order in the legend. Proteins annotated “General – Neuron” have both excitatory and inhibitory neuron annotations.

There were no significant differences in protein expression between SUDEP and non-SUDEP epilepsy cases in any brain region (Fig. 1F-H, fig. S1A-C, Data file S1). Further, a significant correlation of LFQ values for all proteins showed the similarity in protein expression when comparing SUDEP and non-SUDEP epilepsy cases in each brain region by a Pearson’s correlation (p < 0.0001) with the corresponding R^2^ values indicated (fig. S1). Brain cell type specific annotation was evaluated in the 2847 identified proteins, derived from previous methods (*23*), with 19.8% (564/2847) proteins having an annotation while the remaining 80.2% did not and were more ubiquitously expressed or cell type is unknown. Most (78.2%; 502/564) annotated proteins were generally neuronal, with excitatory neuron proteins predominating (48.1%; 271/564) (Fig. 1F-H, Data file S1). Some proteins showed a trend for altered expression in SUDEP cases (p<0.01; table S1–2), but these were not statistically significant at a 5% FDR. Several of these protein changes have been reported in epilepsy animal models and non-epilepsy cases or include proteins encoded by genes in which mutations have been previously linked to epilepsy. However, none of the proteins in table S1–2 have been previously linked to SUDEP pathogenesis. Ermin (ERMN) had the strongest trend for difference in SUDEP with a 2.8-fold decrease in the hippocampal CA1-3 region when comparing SUDEP and non-SUDEP epilepsy cases by MS (fig. S2A). Further, ERMN was detected in more non-SUDEP epilepsy than SUDEP cases by MS, indicating lower abundance of this protein in SUDEP. Validation of the quantitative MS findings with semiquantification of immunohistochemistry (fig. S2B) also showed a decrease of ERMN in SUDEP cases with a 1.3-fold change but was not significant (student’s unpaired t test, p-value = 0.4871). Because ERMN may play a role in myelinogenesis and myelin maintenance, we reviewed the mature oligodendrocyte marker myelin basic protein (MBP) but found no difference between SUDEP and non-SUDEP epilepsy cases in the hippocampal CA1-3 region by MS (fig. S2C).

### Analysis of RNAseq and small RNAseq in low and high-risk SUDEP cases

To determine whether there is a pathological difference in epilepsy cases of low (PGES <50 seconds) and high (PGES ≥50 seconds) risk of SUDEP, RNAseq and small RNAseq analyses were performed on resected surgical frozen hippocampal and temporal cortex tissue. Case histories are summarized in Table 2 and Fig. 2A. A t-SNE (t-distributed stochastic neighbor embedding) plot revealed that anatomical region rather than PGES segregated cases (Fig. 2B). A differential expression analysis comparing the hippocampus of low and high-risk SUDEP cases identified 55 differentially expressed genes: 11 were decreased and 44 were increased in high-risk SUDEP cases (Fig. 2C, Data file S2). Brain cell type specific annotation was evaluated in the 55 differentially expressed genes in the hippocampus, derived from previous methods (*23*), with 14.5% (8/55) of genes having a cell type specific annotation: 4 generally neuronal, 3 excitatory neuron, and 1 inhibitory neuron. The dominant transcripts for the differentially expressed genes in hippocampus were: 37 protein-coding, 15 long non-coding RNAs (lncRNAs), and three awaiting confirmation (Fig. 2D). A Reactome pathway analysis on the 55 significant genes in the hippocampus did not reveal a significant association with any signaling pathways. Several of these genes have been associated with epilepsy human disease and have been studied in animal models, however none of the genes in Table 3 have been linked to SUDEP pathogenesis. The most significantly decreased protein-coding gene in the high-risk SUDEP cases, *GFRA1*, was validated by real time quantitative PCR (RT-qPCR, Table 3, fig. S3). In accordance with the RNAseq analysis, *GFRA1* was decreased 1.7-fold in the high-risk SUDEP cases (Mann-Whitney U test, p-value = 0.0121). In the temporal cortex, one protein-coding gene (*SLC6A5*) with an “undefined” cell type annotation was significantly decreased in the high-risk SUDEP cases, within this small group of cases (Fig. 2E, Data file S3). No genes were differentially expressed in the small RNAseq analyses in the hippocampus and temporal cortex (Data file S4-S5).

**Table 2.**
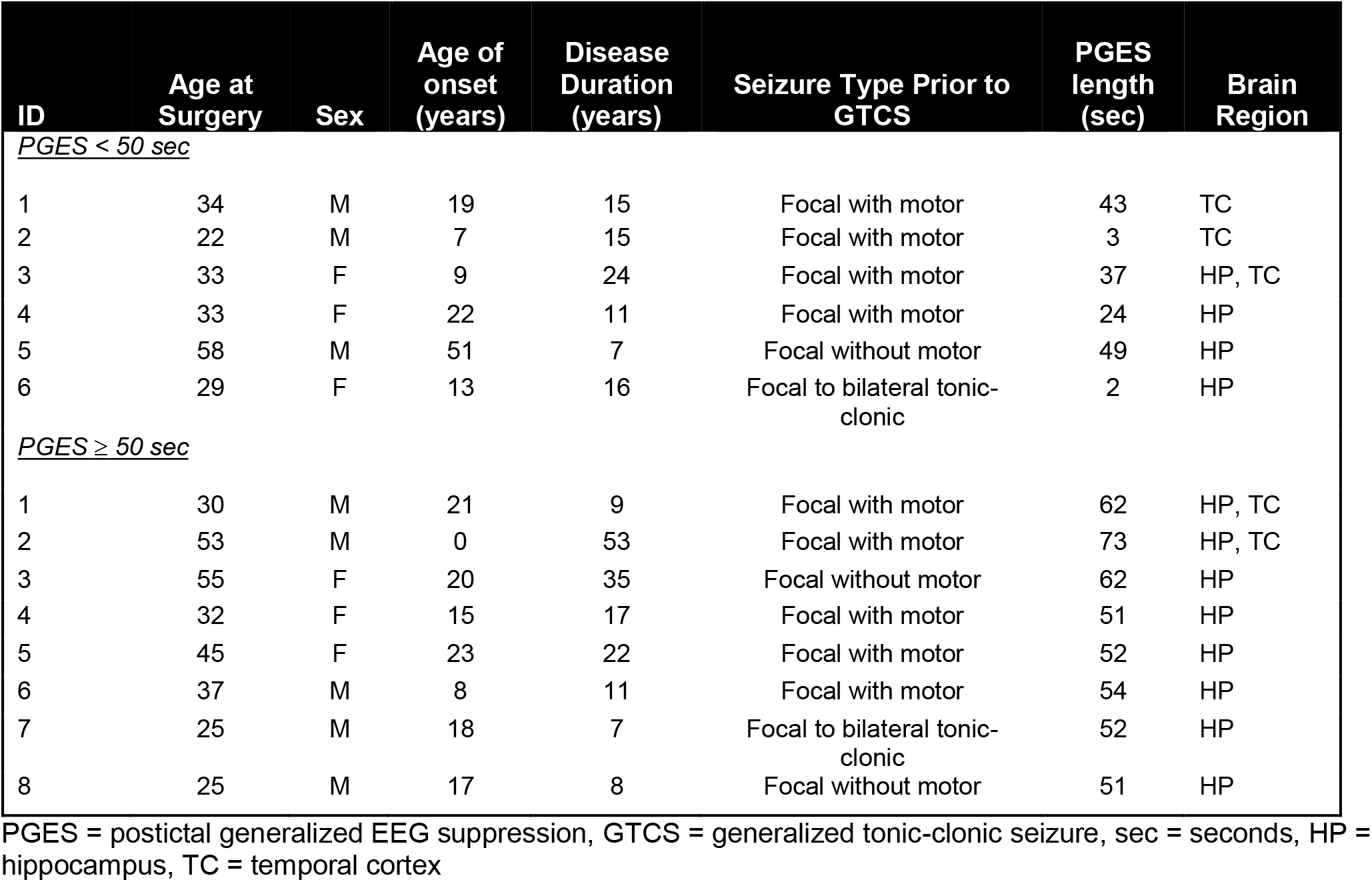
Epilepsy Cases with Low or High-Risk of SUDEP in RNAseq Analyses.

**Table 3.**
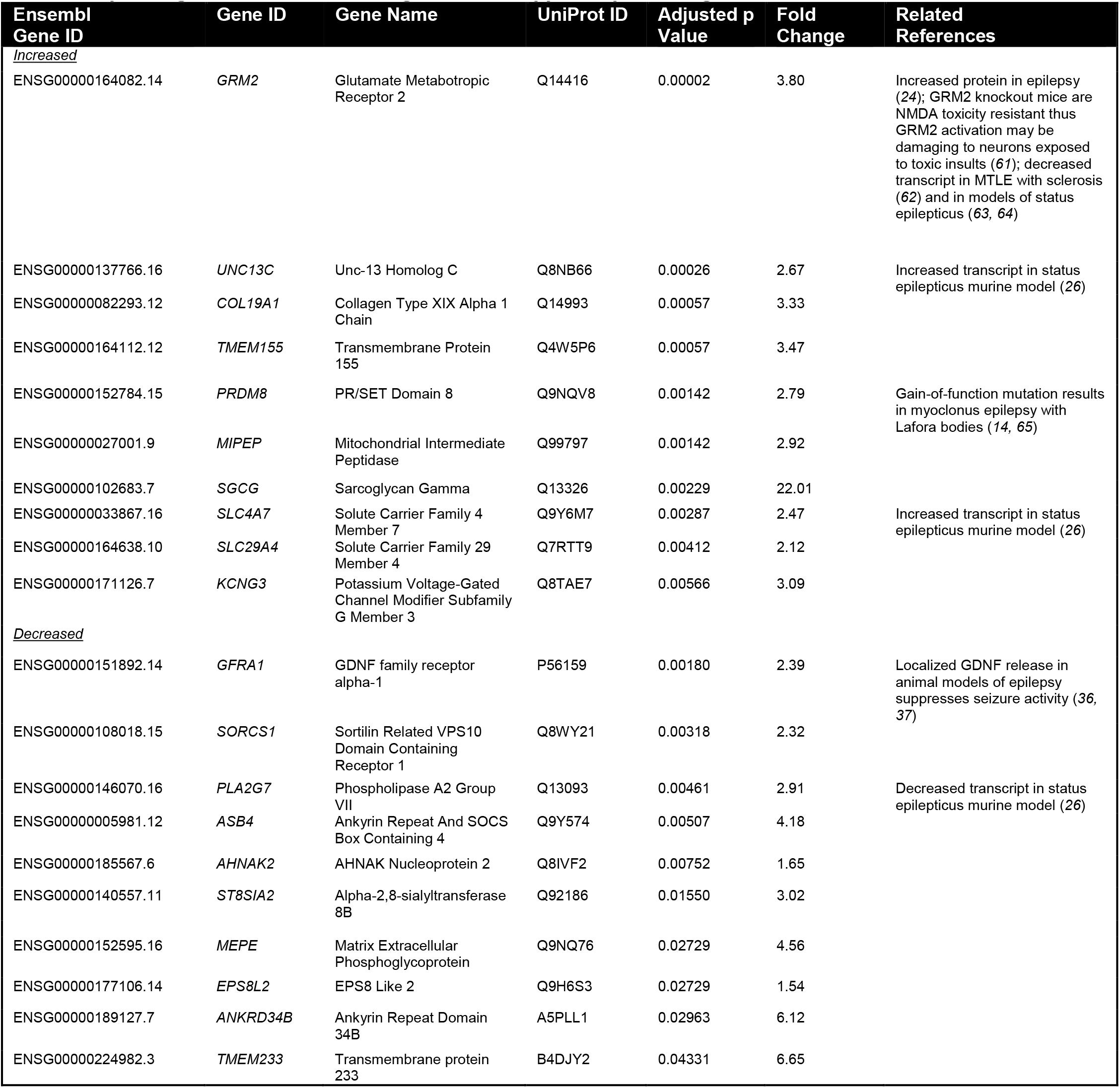
Top 20 Significant Protein-coding Genes in Hippocampus of High vs Low-Risk SUDEP Cases.

**Figure 2.**
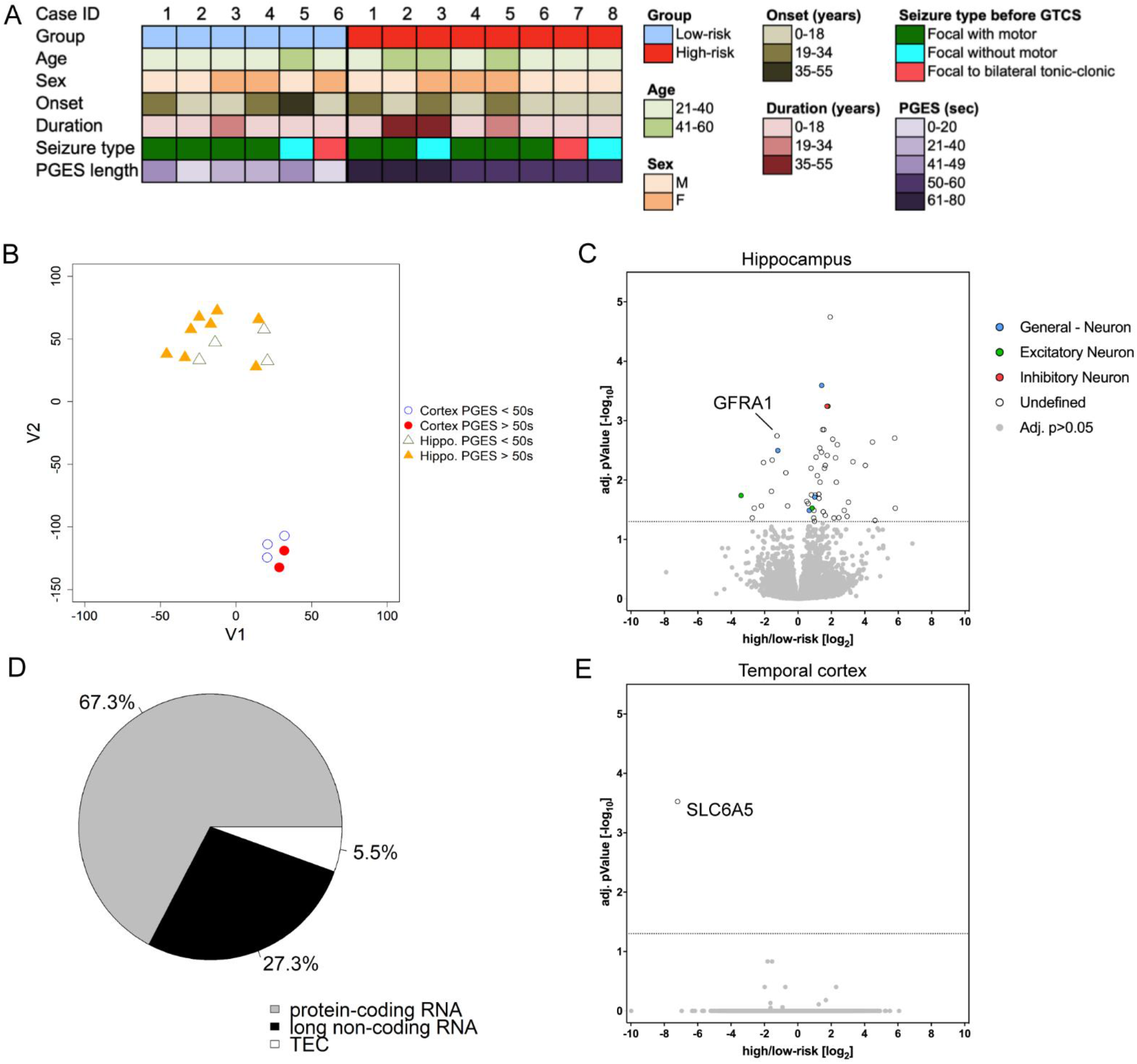
RNAseq of epilepsy cases in the hippocampus and temporal cortex with low and high-risk of SUDEP, as determined by PGES. **A)** Case history summary for low and high-risk SUDEP cases. **B)** The t-SNE (t-distributed stochastic neighbor embedding) plot of RNAseq data shows separation by brain region rather than SUDEP risk status. **C)** A volcano plot shows the results of differential expression analysis of the hippocampus from low-risk (n=4) and high-risk (n=8) SUDEP cases. Eleven genes were decreased and 44 genes were increased in hippocampus of high-risk SUDEP cases. The Wald test identified differentially expressed genes using a Benjamini-Hochberg adjusted p-value <0.05 for significance. Cell type specific gene annotation is included, with the most predominant listed in decreasing order in the legend. Genes annotated “General – Neuron” have both excitatory and inhibitory neuron annotations. **D)** Biotypes of differentially expressed genes are depicted in the hippocampus for high-risk SUDEP compared to low-risk SUDEP cases. Of the 55 differentially expressed genes 67.3% were protein-coding genes, 27.3% were long non-coding RNAs, and 5.5% are yet to be experimentally confirmed (TEC). **E)** A volcano plot shows the results of differential expression analysis in the temporal cortex from low-risk (n=2) and high-risk (n=3) SUDEP cases. One gene was decreased and no genes were increased in the temporal cortex. The Wald test identified differentially expressed genes using a Benjamini-Hochberg adjusted p-value <0.05 for significance.

### Comparison of SUDEP Proteome to High-risk SUDEP Transcriptome

Comparing the 37 differentially expressed protein-coding genes in the RNAseq analyses to the proteomics analyses, only four (GRM2, ERC2, CRTC1, AHNAK2) were detected in the proteomics analyses. Two (GRM2, ERC2) were detected in most cases of the hippocampal CA1-3 region but showed no trend in differential expression for SUDEP cases compared to non-SUDEP epilepsy cases in the proteome.

## Discussion

Our study compared SUDEP or high-risk SUDEP cases to epilepsy controls and revealed no differentially expressed proteins in the hippocampus and frontal cortex and limited transcriptomic changes in the hippocampus and temporal cortex. Thus, the proteome in SUDEP and transcriptome in high-risk SUDEP largely reflects other epilepsy cases, consistent with the diverse spectrum of syndromes and severities associated with SUDEP (*5*). In the hippocampus, the few differentially expressed genes identified in high-risk SUDEP cases included a high proportion of lncRNAs (15/55, 27%). Given that we detect robust proteome (*24*) and transcriptome (*25*) differences in the hippocampus and cortex with similar group sizes for epilepsy and non-epilepsy control cases, our data in this study suggest that these brain regions are not especially or uniquely affected in SUDEP.

To validate the label-free quantitative MS findings, immunohistochemistry was used to confirm changes in ermin (ERMN) expression, as this protein had the strongest trend for difference in SUDEP. Immunohistochemistry results corroborated a trend in a decreased fold change of ERMN in the hippocampal CA1-3 region of SUDEP cases when compared to non-SUDEP epilepsy cases, although this similarly was not significant. Further, *ERMN* was not significantly altered in the current RNAseq study or in our previous proteomics analyses between non-SUDEP epilepsy and controls (*24*). However, in our previous RNAseq study between MTLE and non-epilepsy controls, *ERMN* was decreased (*25*) and is reportedly decreased in a murine model of status epilepticus (*26*). Expressed by oligodendrocytes, ERMN regulates cytoskeleton arrangement during myelinogenesis and myelin sheath maintenance (*27*). Myelin damage may occur after prolonged seizures and its loss may promote further seizure activity (*28–30*). We found that the mature oligodendrocyte marker myelin basic protein (MBP) is decreased in epilepsy cases compared to non-epilepsy control cases (*24*), and it is decreased in the hippocampus of an animal model of epilepsy (*31*). However, we found no further decrease in MBP expression in SUDEP or high-risk SUDEP cases when compared to controls in this study, nor was *MBP* different in our recent RNAseq analysis between MTLE and non-epilepsy controls (*25*). Overall, ERMN is significantly decreased in surgical MTLE versus non-epilepsy controls at the transcriptomic level (*25*) and trending to decrease in protein expression of SUDEP versus non-SUDEP epilepsy, indicating that ERMN may be decreased in response to the elevated seizure activity that may be seen in refractory epilepsy that requires surgery and in some cases of SUDEP. The impact on myelination, as measured by MBP, is only apparent in these cases for protein expression rather than gene expression in epilepsy versus non-epilepsy controls with no further decrease in SUDEP. Thus, further investigation should assess the potential role of ERMN in epilepsy and SUDEP, and whether reduced ERMN may reflect the severity of pathology resulting from seizure burden in some SUDEP cases.

The RNAseq and small RNAseq analyses showed moderate changes in the hippocampus and minimal differences in the temporal cortex in high-risk compared to low-risk SUDEP MTLE cases. Interestingly, 15/55 differentially expressed genes in the hippocampus were lncRNAs. LncRNAs are an understudied transcriptomic component implicated in many neurological disorders (*32, 33*), but few studies have been done regarding their role in epilepsy or SUDEP (*34*). Among the protein-coding genes differentially expressed in the hippocampus, *GFRA1* (GDNF Family Receptor Alpha 1) was the most decreased. GDNF (glial cell-derived neurotrophic factor) binds to GFRA1 and plays a role in neuronal survival and differentiation, including that of GABAergic interneurons (*35*). Localized release of GDNF in the hippocampus of an animal model of epilepsy suppresses seizure activity (*36, 37*). Thus, decreased *GFRA1* may reflect a change in cell survival or result in reduced GDNF mediated seizure suppression in high-risk SUDEP cases. Of the top 20 differentially expressed genes (Table 3), *SGCG* (Sarcoglycan Gamma) had the largest change at a 22.0-fold increase (adjusted p=0.0023) in the high-risk SUDEP cases. SGCG is expressed in the cerebrovascular system and may localize to vascular smooth muscle cells, potentially involved in membrane contractility, stabilization, and signaling in the associated dystrophin complex affecting neurovascular coupling (*38*). Its neural role is unknown, but aberrant cerebrovascular organization occurs in MTLE (*39*). Additional studies are needed to determine how the altered levels of some protein-coding genes and lncRNAs we identified may impact mechanisms related to SUDEP risk.

Protein expression in the brain has rarely been studied in human SUDEP. Hippocampal HSP70 positive neurons are reportedly increased in post-mortem SUDEP cases when compared to non-SUDEP epilepsy cases, but similar to surgical epilepsy cases, suggesting this is likely related to ante-mortem neuronal injury perhaps due to a terminal seizure in SUDEP cases (*40*). HSP70 expression was similar in both the proteomic and RNAseq analyses among our cases. Another immunohistochemistry study found few differences in the hippocampus, amygdala, and medulla of post-mortem SUDEP compared to non-SUDEP epilepsy and non-epilepsy control cases with minimal significant changes reported for several markers of inflammation (CD163, HLA-DR, GFAP), compromised blood brain barrier (IgG, albumin), and HIF-1α, a transcriptional regulator of cellular responses to hypoxia (*20*). We found increased GFAP in the hippocampus of three epilepsy cases (3/26, 11.5%); two had gliosis independent of SUDEP status. GFAP was not increased in most non-SUDEP epilepsy cases when compared to non-epilepsy control cases (*24*), but it was increased in the hippocampus of one (1/14, 7.1%) epilepsy case with hippocampal gliosis. Increased GFAP occurs in some epilepsy cases and after prolonged seizures in rodent models (*41, 42*). Further, *GFAP* was not altered in MTLE cases with high-risk of SUDEP in the current RNAseq analysis, but this gene was significantly increased in the hippocampus of MTLE cases compared to non-epilepsy controls (*25*).

Our study had some limitations. The LCM derived label-free quantitative MS allows for detection of localized protein changes that would not be possible using bulk homogenate, however this technique detects a lower quantity of membrane proteins that are relatively insoluble with this method. Thus, we may not detect differential expression of some membrane proteins, although downstream signaling pathways reflecting their functional activity may be identified. Additional limitations include the heterogeneity of epilepsies, seizure types, and neuropathology due to case availability, and further reinforces the importance of banking various brain tissue samples from SUDEP cases. Our study was powered to identify proteomic differences across the representative SUDEP group rather than epilepsy-subgroups. Potential pathogenic gene variants were not assessed in our patients. Our proteomics analyses were based on NASR referrals, skewed by major referral sources: the San Diego Medical Examiner Office (mainly low socioeconomic white and Hispanic patients) and direct referrals (mainly high socioeconomic white patients). For the RNAseq analyses, surgical cases had treatment-resistant MTLE. PGES duration as a biomarker of SUDEP risk has not been validated, can vary within the same patient for different seizures, and the number video EEG-recorded GTCS in each patient was limited (*10, 43–45*). Thus, group differences may reflect sampling bias. Further, the number of cases used for the RNAseq temporal cortex analyses was low. Finally, further investigation is needed in brain regions implicated in SUDEP, including the brainstem, as it modulates autonomic functions and it has been suggested that seizure-induced postictal depression of arousal, respiratory, and cardiac function may occur in SUDEP (*46–48*).

In summary, in contrast to robust differences we found in proteomic and RNAseq analyses between epilepsy and non-epilepsy cases (*24, 25*), there were no differences detected in the proteomic analyses of autopsy tissue from SUDEP and non-SUDEP epilepsy cases and limited transcriptomic differences comparing surgical tissue from low and high-risk SUDEP cases in the brain regions analyzed, consistent with the diverse epilepsy syndromes and comorbidities associated with SUDEP and indicating that epilepsy subtypes and additional brain regions should be examined further.

## Materials and Methods

### Human Brain Tissue for Proteomics

Post-mortem brain tissue from epilepsy patients who died from SUDEP or other causes was obtained through the North American SUDEP Registry (NASR) with approval by the New York University School of Medicine Institutional Review Board (IRB). Causes of death were classified (OD, DF) into non-SUDEP epilepsy and SUDEP (definite SUDEP, definite SUDEP plus, and probable SUDEP) (*4, 5*). Lifetime GTCS history was determined from interviews and medical records, representing the best estimate for each case and as described previously (*5*). After neuropathological review (TW, AF), brain tissue was processed into formalin fixed paraffin embedded (FFPE) blocks and sections were stained with luxol fast blue counterstained with hematoxylin & eosin (LFB/H&E). Archival time for brain tissue storage in formalin was less than or equal to three years. Clinical and neuropathologic data on the 14 non-SUDEP epilepsy and 12 SUDEP cases are summarized in Table 1. Group sizes were determined based on the number of cases with significant findings as previously reported (*49–51*), including our earlier studies in epilepsy cases with similar methods (*24, 25*).

### Laser Capture Microdissection for Proteomics

FFPE brain tissue blocks containing either hippocampus (lateral geniculate nucleus level) (*52, 53*) or superior frontal gyrus were sectioned at 8 μm and collected onto laser capture microdissection (LCM) compatible PET slides (Leica). Sections were stained with cresyl violet to localize regions of interest for LCM (*54*) and air dried overnight in a loosely closed container. LCM was used to individually microdissect 10 mm^2^ from the hippocampal CA1-3 region and superior frontal cortex (layers I-IV), and 4 mm^2^ from the hippocampal dentate gyrus into LC-MS grade water (Thermo Scientific). Microdissected samples were centrifuged for 2 minutes at 14,000*g* and stored at −80°C. LCM was performed at 5X magnification with a LMD6500 microscope equipped with a UV laser (Leica).

### Label-free quantitative MS Proteomics

Label-free quantitative MS assessed differential protein expression, as described previously (*24*). Samples were thawed at room temperature, resuspended (in 200 μl of 100 mM ammonium bicarbonate, 20% acetonitrile solution) and an additional 50 μl was added to the cap and spun down. This step was repeated three times to thoroughly wash the cap. Samples were then incubated at 95°C for 1 hour, followed by incubation at 65°C for 2 hours. Reduction was performed with DTT at 57°C for 1 hour (2 μl of 0.2 M) followed by alkylation with iodoacetamide at room temperature in the dark for 45 minutes (2 μl of 0.5 M). Samples were digested with shaking overnight at room temperature in 300 ng of sequencing grade modified trypsin (Promega). Samples were then acidified with trifluoroacetic acid (TFA) to pH ~2. For protein extraction, 15 μl of R2 20 μm POROS beads slurry (Life Technologies Corporation) were added to the sample followed by a 3-hour incubation at 4°C on a shaker. The beads were then loaded onto equilibrated C18 ZipTips (Millipore) using a microcentrifuge for 30 seconds at 3000 rpm. Sample vials and caps were rinsed three times with 0.1% TFA and each rinse was added to its corresponding ZipTips followed by microcentrifugation. The extracted POROS beads were washed with 0.5% acetic acid and the peptides were eluted with 40% acetonitrile in 0.5% acetic acid. The second elution step used 80% acetonitrile in 0.5% acetic acid. The organic solvent was removed using a SpeedVac concentrator and the sample reconstituted in 0.5% acetic acid. For proteomics analysis, samples were individually analyzed using the autosampler of an EASY-nLC 1200 Liquid Chromatograph (Thermo Scientific). The peptides were gradient eluted from the column directly to Orbitrap Fusion Lumos mass spectrometer (Thermo Scientific) using a 165min gradient (Thermo Scientific, Solvent A consisting of 2% acetonitrile, 0.5% acetic acid and Solvent B of 80% acetonitrile, 0.5% acetic acid). High resolution full mass spectrometry (MS) spectra were acquired with a resolution of 240,000, an automatic gain control (AGC) target of 1e6, with a maximum ion time of 50 ms, and scan range of 400-1500 m/z. After each full MS, MS/MS HCD spectra were acquired in the ion trap. All MS/MS spectra were collected using specific instrument parameters: Ion trap scan rate Rapid, AGC target of 2e4, maximum ion time of 18 ms, one microscan, 2 m/z isolation window, fixed first mass of 110 m/z, and NCE of 32.

### Proteomics Computational Analysis

MS data were analyzed using MaxQuant software (*55*) version 1.6.3.4 and searched against the SwissProt subset of the human UniProt database (http://www.uniprot.org/) containing 20,421 entries. Database search used Andromeda (*56*) integrated in MaxQuant environment, including 248 common laboratory contaminants as well as reversed versions of all sequences. For searching, the enzyme specificity was set to trypsin with the maximum number of missed cleavages of 2. The precursor mass tolerance was 20 ppm for the first search used for non-linear mass re-calibration and then to 6 ppm for the main search. Oxidation of methionine was searched as a variable modification; carbamidomethylation of cysteines was searched as a fixed modification. The false discovery rate (FDR) for peptide, protein, and site identification was set to 1%, the minimum peptide length was set to 6. To transfer identifications across different runs, the ‘match between runs’ option in MaxQuant was enabled with a retention time (RT) window of 0.7 minutes (with a window of 20 minutes for initial RT alignment). Protein quantification was performed with the built-in MaxQuant LFQ algorithm with the following settings: minimum ratio count of 2, “fastLFQ” option enabled, minimum/average number of neighbors 3/6. LFQ normalization was performed individually within groups of raw files corresponding to the different brain regions (dentate gyrus, hippocampal CA1-3 region, and frontal cortex) due to expected large variability in their proteome composition.

Subsequent data analysis steps used either Perseus (http://www.perseus-framework.org/) or R environment for statistical computing and graphics (http://www.r-project.org/).

### Proteomics Statistical Analyses

The protein expression matrix (n=4,129) was filtered to contain only proteins that were quantified in ≥ 8 replicates in at least 1 condition (SUDEP or epilepsy) in any brain region (n=2,847). For Principal Component Analysis (PCA), missing values were imputed from the normal distribution with a width of 0.3 and downshift of 1.8 (relative to measured protein intensity distribution). An unpaired two-tailed t test was performed for PCA1 in each brain region to determine significance of separation in the SUDEP and non-SUDEP epilepsy cases. All other analyses were done using nonimputed data. A Student’s two sample t-test was used to detect significant changes in protein expression between conditions. Thresholds set for p value were adjusted to provide a false discovery rate (FDR) below 5% (permutation-based FDR from 250 data randomizations). Cell type specific annotations were included in the Data file S1 and on volcano plots in Fig. 1F-H, derived from previous data (*23*). Annotations were included when a protein had only one associated cell type after removing cerebellar annotations and when the annotation included more than one associated cell type (both excitatory and inhibitory neuron annotations) and were thus assigned a general neuron annotation, for a total of 1066 possible annotations.

### Proteomics Correlation

Average LFQ values for each protein group of the SUDEP or non-SUDEP epilepsy groups were evaluated in each brain region to determine the correlation in protein abundance. A Pearson’s correlation was performed for proteins detected in both SUDEP and non-SUDEP epilepsy cases for each brain region, with 2715 proteins for hippocampal CA1-3, 2464 proteins for dentate gyrus, and 2695 proteins for the frontal cortex.

### Immunohistochemistry

Immunohistochemistry was performed to validate the identified protein of interest, ermin (ERMN) as previously described (*24, 57*). Briefly, FFPE sections (8 μm) were deparaffinized and rehydrated through a series of xylenes and ethanol dilutions. Heat-induced antigen retrieval was performed with 10 mM sodium citrate, 0.05% triton-x 100; pH6. Blocking with 10% normal donkey serum was followed by ERMN primary antibody (1:200, Sigma HPA038295) overnight at 4°C. Sections were incubated with donkey anti-rabbit Alexa-Fluor 647 secondary antibody (1:500, Thermofisher Invitrogen) and coverslipped.

### Image semiquantitative analysis

Whole slide scanning was performed at 20X magnification with a NanoZoomer HT2 (Hamamatsu) microscope using the same settings for each slide. One image containing the hippocampal CA1-3 region was collected for each case, 11 non-SUDEP epilepsy and 11 SUDEP cases. Images were analyzed in Fiji ImageJ to compare the amount of ERMN in SUDEP and non-SUDEP epilepsy cases. The same binary threshold was used for all images to determine the number of ERMN positive pixels in each image, which was reported as a percentage of the total image area. An unpaired t-test was performed for statistical analysis; p-value <0.05 was considered significant.

Confocal imaging was used to collect representative images of ERMN immunohistochemistry, using a Zeiss LSM800 confocal microscope with the same settings on each slide with a Plan-Apochromat 20X/0.8 M27 objective and a pinhole of 38 μm.

### RNA-sequencing datasets

Small RNA-sequencing (small RNAseq) and RNA-sequencing (RNAseq) data sets were retrieved form the European Genome-phenome Archive (accession number: EGAS00001003922) (*25*). Small RNAseq and RNAseq data was retrieved for 6 cases with PGES < 50 sec, indicating a potential low-risk for SUDEP, and 8 cases with PGES ≥ 50 sec, indicating a potential high-risk for SUDEP as previously described (*8, 9*). Table 2 summarizes the clinical characteristics of these cases. PGES occurrence and duration was assessed by two epileptologists (CS, RT).

### Bioinformatic analysis of RNAseq data

Bioinformatic analysis was performed as described previously (*25*). Briefly, read quality was assessed using FastQC v0.11.2 software produced by the Babraham Institute (Babraham, Cambridgeshire, UK) and Trimmomatic v0.36 filtered reads of low quality. Low quality leading and trailing bases were removed from each read, a sliding window trim using a window of 4 and a phred33 score threshold of 20 was used to assess the quality of the body of the read. Any read <75 nucleotides in length was excluded from the analysis. Only reads with forward and reverse elements were included in the downstream analysis. Reads were aligned to the human reference genome, GRCh38 using TopHat2 v2.0.13 default settings. The featureCounts program from the Subread package calculated the number of reads aligned to each gene using Gencode v27. Library normalization and differential expression testing was carried out using the R package *DESeq2*. The Wald test identified differentially expressed genes using a Benjamini-Hochberg adjusted p-value <0.05 for significance. Cell type specific annotations were included in the Data file S2, Data file S3, and on volcano plots in Fig. 2C,2E, derived from previous data (*23*). Annotations were included when a protein had only one associated cell type after removing cerebellar annotations and when the annotation included more than one associated cell type (both excitatory and inhibitory neuron annotations) and were thus assigned a general neuron annotation, for a total of 1066 possible annotations.

A Reactome pathway enrichment analysis was performed using the R package *ReactomePA*. The differentially expressed genes from the RNAseq differential expression analysis were put into R and tested for over-representation of enriched Reactome pathways using hypergeometric testing. Pathways with a Benjamini-Hochberg corrected p-value <0.05 were considered significantly enriched.

### Bioinformatic analysis of small RNAseq data

Bioinformatic analysis of the small RNAseq data was performed as described previously (*25*). Briefly, read quality was assessed using FastQC v0.11.2 software produced by the Babraham Institute (Babraham, Cambridgeshire, UK) and Trimmomatic v0.36 was used to filter reads of low quality. Low quality leading and trailing bases were removed from each read, a sliding window trim using a window of 4 and a phred33 score threshold of 15 assessed the quality of the body of the read. Reads <17 nucleotides were excluded from the analysis. Only reads with forward and reverse elements were included in the downstream analysis. Reads were aligned to the human reference genome, GRCh38 using Bowtie2 v2.3.2, no mismatches between the seed sequence and the reference genome were allowed, and reads were allowed to align a maximum of ten times. Using the featureCounts program from the Subread package the number of reads that aligned to miRNAs in accordance to miRbase21 (www.mirbase.org) and other short RNA species extracted from Gencode v27 were calculated. Library normalization and differential expression testing was carried out using the R package *DESeq2*. The Wald test identified differentially expressed genes with a Benjamini-Hochberg adjusted p-value <0.05 considered significant.

### RNAseq validation by qPCR

The gene expression of GDNF Family Receptor Alpha 1 (*GFRA1*) was assessed in the same cohort of samples used in the RNAseq analysis for which sufficient RNA remained (PGES < 50s, n=4, PGES ≥ 50s, n=7). PCR primers based on the reported cDNA sequences were designed using the NCBI primer design tool (*58*). The sequences for the forward and the reverse primers of GFRA1 were 5’-TCT TCC AGC CGC AGA AGA AC-3’ and 5’-AAC AGT GGG GAC AAA CTG GG-3’ respectively. 700 ng of total RNA was reverse transcribed into cDNA using oligodT primers. For each qPCR reaction, a mastermix was prepared as follows: 1 μl cDNA, 2.5 μl of 2x SensiFAST SYBR Green Reaction Mix (Bioline Inc, Taunton, MA, USA), 0.2 μM of both reverse and forward primers and the PCRs were run on a Roche Lightcycler 480 thermocycler (Roche Applied Science, Basel, Switzerland). Each sample and primer pair was run in triplicates. Data quantification was performed using the software LinRegPCR in which linear regression on the Log (fluorescence) per cycle number data determined amplification efficiency per sample (*59, 60*). The starting concentration of each specific product was divided by the geometric mean of the starting concentration of the reference genes, Eukaryotic Translation Elongation Factor 1 Alpha 1 (*EEF1A1*) and Chromosome 1 Open Reading Frame 43 (*C1orf43*). This ratio was compared between the two groups (Mann-Whitney U test); p <0.05 was considered significant.

## General

The authors wish to thank the participating families and clinicians for their involvement with the North American SUDEP Registry (NASR).

## Author contributions

O.D. and E.A. conceived the study. D.L., J.D.M., E.D., and G.P. designed and performed the experiments and data analysis. A.F., D.F., C.V., T.W., and O.D. were involved in tissue sample and/or clinical data acquisition for proteomics analyses. E.K., S.N., M.A., and B.U. performed initial proteomics analyses. J.C.B., R.T., S.I., J.J.A., M.T., E.A., O.D., C.S., B.D. contributed to the acquisition of the surgical tissue samples and/or clinical data for RNAseq analyses. J.D.M. performed the RNAseq bioinformatics analysis, with the support of B.J.C. and M.J. All wet-laboratory experiments were performed by D.L., J.D.M., G.P., J.D.M., E.A.vV., and J.J.A. C.S. and R.T. evaluated PGES. D.L., G.P., E.D., J.D.M., E.A.vV., D.F., S.D., E.A., and O.D. helped with data interpretation and the writing of the manuscript. All authors read, revised, and approved the final manuscript.

## Data and materials availability

All data needed to evaluate the conclusions in the paper are present in the paper and/or the Supplementary Materials. Additional data related to this paper may be requested from the authors.

## Supplementary Materials

Data file S1. MS raw data for Epilepsy and SUDEP Cases

Data file S2. RNAseq raw data from hippocampus in Low and High-Risk SUDEP

Data file S3. RNAseq raw data from temporal cortex in Low and High-Risk SUDEP

Data file S4. Small RNAseq raw data from hippocampus in Low and High-Risk SUDEP

Data file S5. Small RNAseq raw data from temporal cortex in Low and High-Risk SUDEP

---

**Figure S1.**
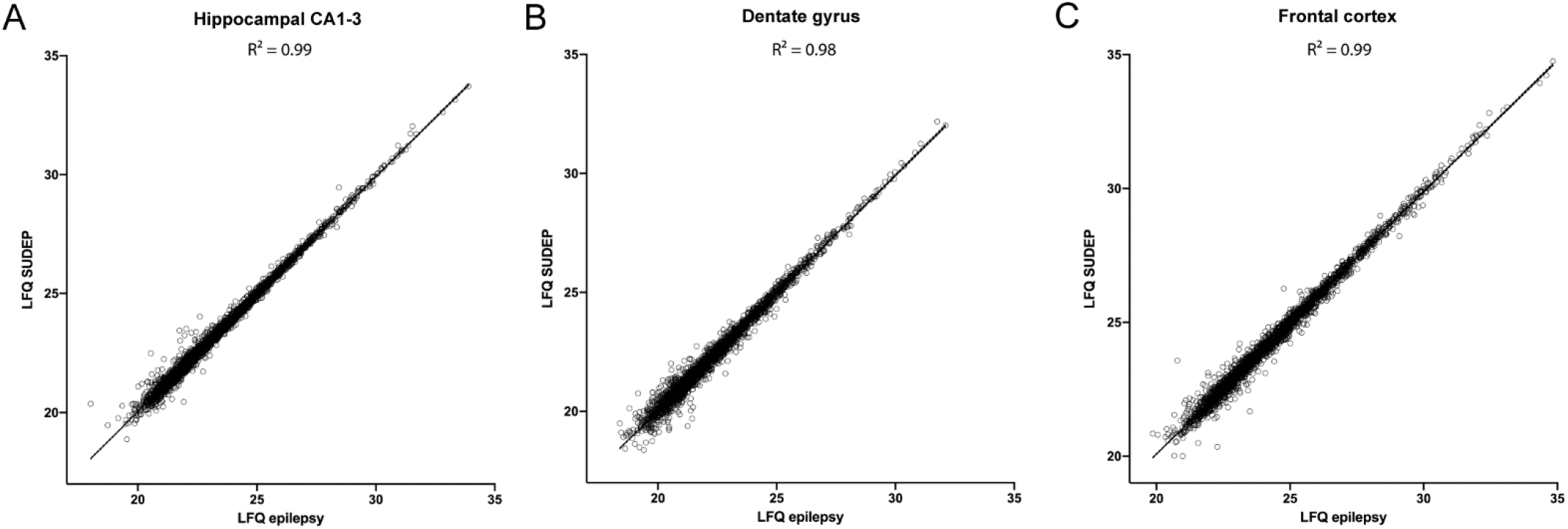
Correlation of proteins detected in each brain region of SUDEP and non-SUDEP epilepsy cases. **A-C)** A correlation of LFQ values is significant in each brain region by a Pearson’s correlation (p < 0.0001) with the corresponding R^2^ value indicated, showing the similarities in protein expression when comparing SUDEP and non-SUDEP epilepsy cases. Confidence intervals of 95% are represented for the linear regression.

**Figure S2.**
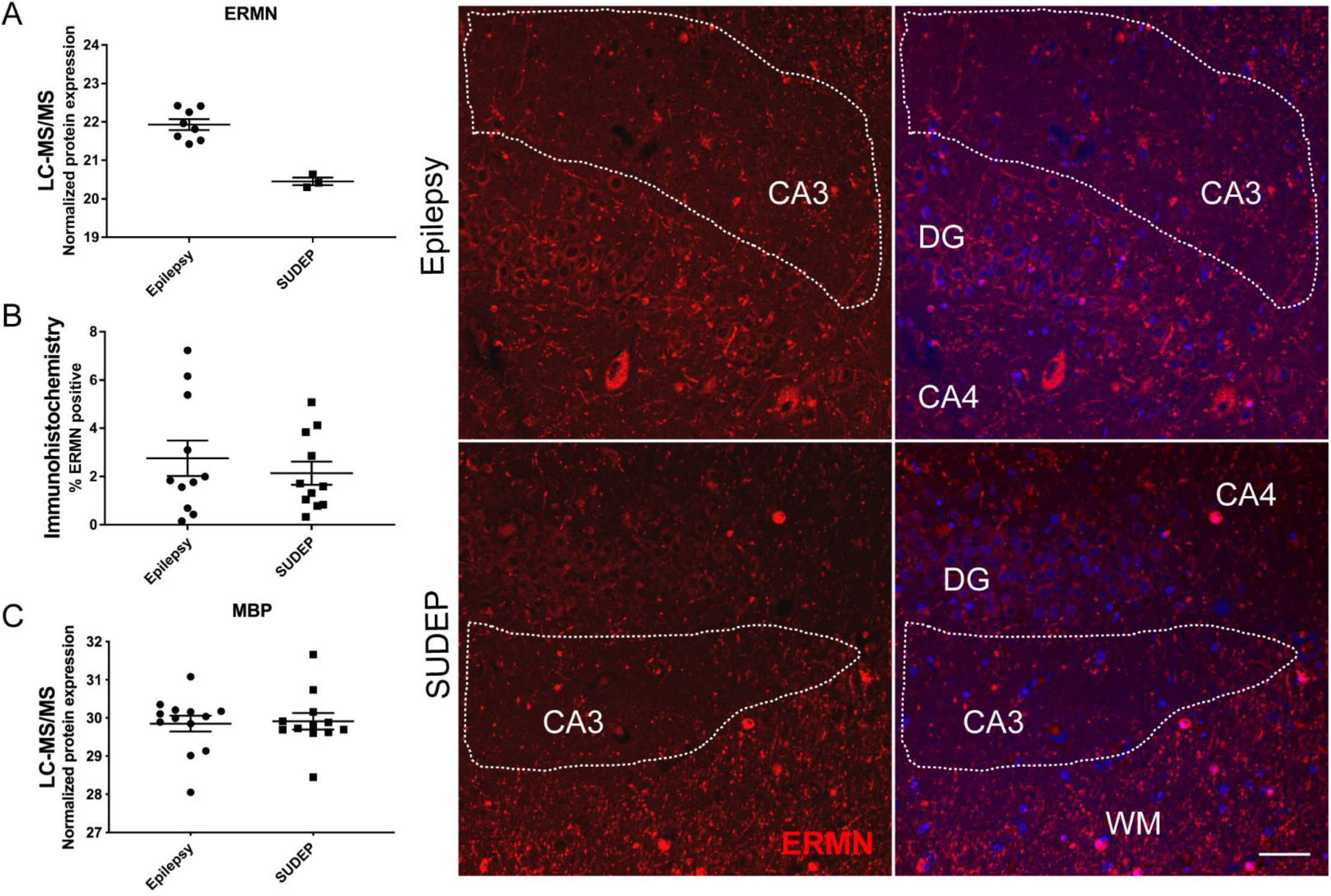
ERMN and MBP protein expression in the hippocampal CA1-3 region of SUDEP and non-SUDEP epilepsy cases. **A)** Quantification from MS of ERMN was evaluated in non-SUDEP epilepsy cases (n=14) and SUDEP cases (n=12) in the hippocampal CA1-3 region. ERMN was detected in n=8 non-SUDEP epilepsy and only n=3 SUDEP cases. As determined by a student’s two tailed t-test with permutation correction at a 5% FDR, ERMN is not significantly different. **B)** Immunohistochemistry of ERMN in the hippocampal CA1-3 region of non-SUDEP epilepsy cases (n=11) and SUDEP cases (n=11) shows by semiquantitative analysis that ERMN expression follows the same trend seen in MS (student’s unpaired t test, p-value = 0.4871). ERMN is present at a low level in the CA3 region adjacent to the dentate gyrus (DG), as well as in neighboring white matter (WM) at higher levels. Scale bar indicates 50 μm. **C)** MBP quantification by MS in the hippocampal CA1-3 region shows that there is no significant difference between SUDEP and non-SUDEP epilepsy cases as determined by a student’s two tailed t-test (p = 0.8380) with permutation correction at a 5% FDR. Error bars indicate SEM.

**Figure S3.**
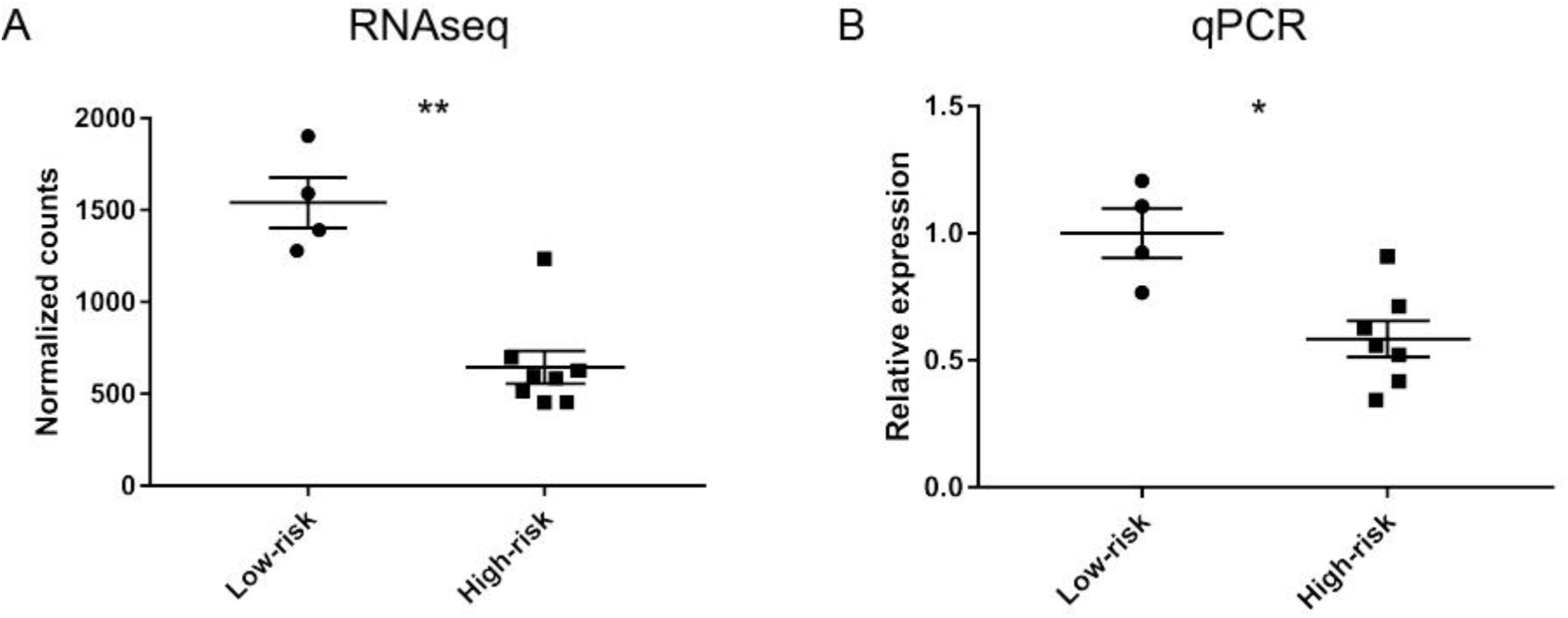
*GFRA1* gene expression in the hippocampus of low and high-risk SUDEP cases by RNAseq and qPCR analysis. **A)** RNAseq analysis shows that *GFRA1* gene expression is significantly decreased (2.386-fold, adjusted p-value = 0.0018) in high-risk SUDEP cases (n=8) when compared to low-risk SUDEP cases (n=4) in the hippocampus. The Wald test identified differential expression using a Benjamini-Hochberg adjusted p-value <0.05 for significance. **B)** Real time quantitative PCR (RT-qPCR) validated the RNAseq findings, with a significant decrease in *GFRA1* of 1.71-fold in high-risk SUDEP cases (n=7) when compared to low-risk SUDEP cases (n=4) in the hippocampus (Mann-Whitney U test, p-value = 0.0121). A p-value <0.05 was considered significant. Mean is indicated ± SEM.

**Table S1.**
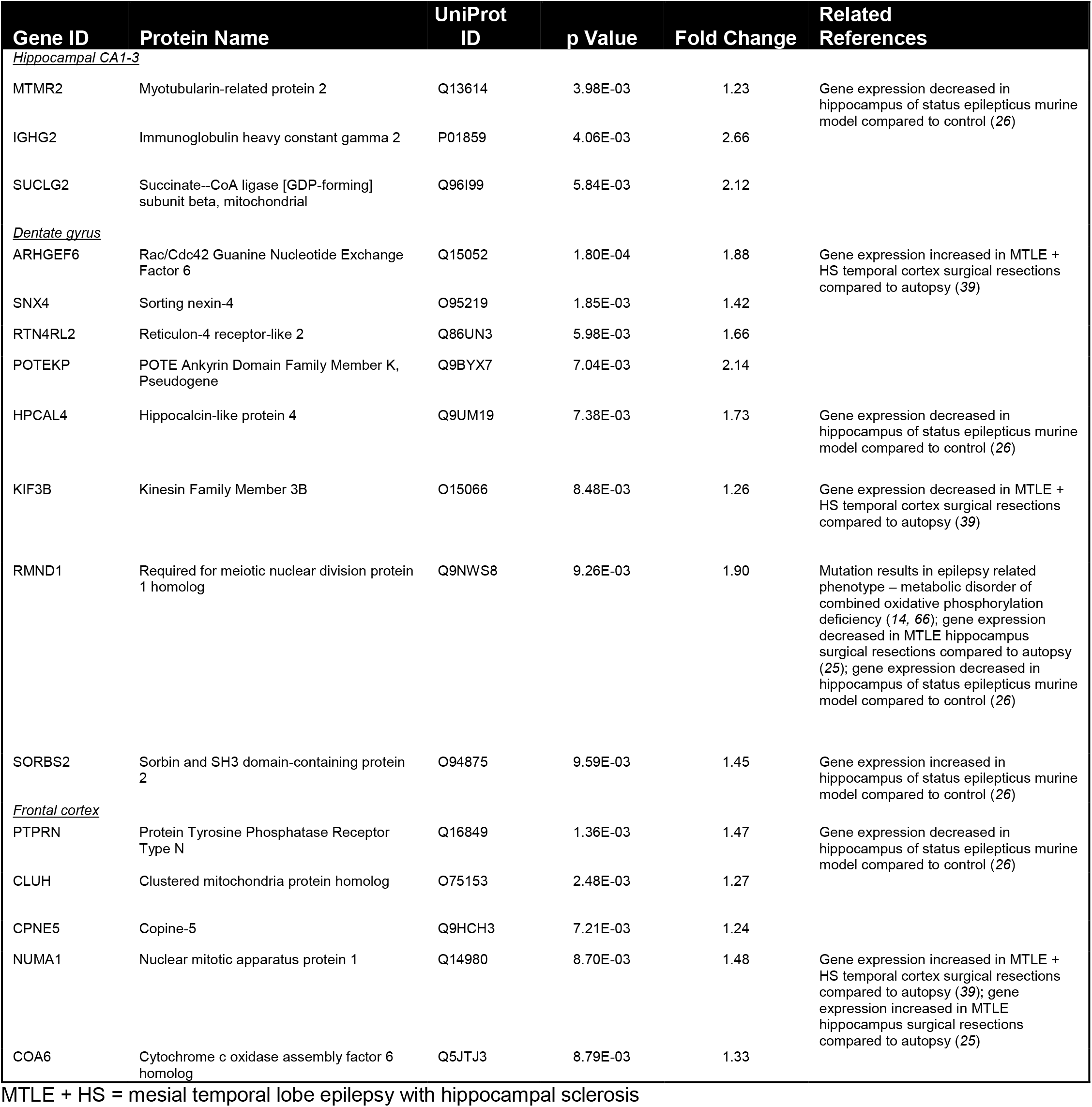
Increased Proteins at p<0.01 in SUDEP versus non-SUDEP Epilepsy.

**Table S2.** Decreased Proteins at p<0.01 in SUDEP versus non-SUDEP Epilepsy.

| Gene ID | Protein Name | UniProt ID | p Value | Fold Change | Related References |
| --- | --- | --- | --- | --- | --- |
| <u>Hippocampal CA1-3</u> |  |  |  |  |  |
| ERMN | Ermin | Q8TAM6 | 1.86E-04 | 2.79 | Gene expression decreased in MTLE hippocampus surgical resections compared to autopsy (25); gene expression decreased in hippocampus of status epilepticus murine model compared to control (26) |
| DDI2 | DNA Damage Inducible 1 Homolog 2 | Q5TDH0 | 5.60E-03 | 1.42 | Gene expression increased in MTLE hippocampus surgical resections compared to autopsy (25); gene expression increased in corpora quadrigemina of audiogenic seizure rat model compared to control (67); gene expression increased in hippocampus of status epilepticus murine model compared to control (26) |
| ARF6 | ADP-ribosylation factor 6 | P62330 | 7.97E-03 | 1.44 | Gene expression decreased in MTLE hippocampus surgical resections compared to autopsy (25); knockdown in the dentate gyrus increases susceptibility to seizures in a mouse model (68) |
| <u>Dentate gyrus</u> |  |  |  |  |  |
| NUTF2 | Nuclear transport factor 2 | P61970 | 3.95E-03 | 1.59 | Gene expression increased in MTLE + HS temporal cortex surgical resections compared to autopsy (39) |
| NDUFB6 | NADH dehydrogenase [ubiquinone] 1 beta subcomplex subunit 6 | O95139 | 5.72E-03 | 1.77 |  |
| TGM2 | Transglutaminase 2 | P21980 | 5.96E-03 | 1.36 | Protein increased early in hippocampus of status epilepticus rat model (69, 70) |
| FAM49B | Family With Sequence Similarity 49 Member B | Q9NUQ9 | 8.45E-03 | 1.46 | Gene expression decreased in MTLE + HS temporal cortex surgical resections compared to autopsy (39); gene expression decreased in MTLE hippocampus surgical resections compared to autopsy (25) |
| SNX2 | Sorting nexin-2 | O60749 | 9.28E-03 | 1.31 | Gene expression increased in MTLE + HS temporal cortex surgical resections compared to autopsy (39); gene expression decreased in hippocampus of status epilepticus murine model compared to control (26) |
| <u>Frontal cortex</u> |  |  |  |  |  |
| COX7A2L | Cytochrome c oxidase subunit 7A-related protein, mitochondrial | O14548 | 3.72E-04 | 1.64 | Gene expression decreased in MTLE + HS temporal cortex surgical resections compared to autopsy (39); gene expression decreased in MTLE hippocampus surgical resections compared to autopsy (25); gene expression decreased in hippocampus of status epilepticus murine model compared to control (26) |
| ADGRB3 | Adhesion G protein-coupled receptor B3 | O60242 | 6.29E-04 | 1.47 |  |
| VPS52 | Vacuolar protein sorting-associated protein 52 homolog | Q8N1B4 | 6.52E-03 | 1.33 | Gene expression increased in MTLE + HS temporal cortex surgical resections compared to autopsy (39) |
| CORO7 | Coronin-7 | P57737 | 6.98E-03 | 1.31 | Gene expression decreased in MTLE hippocampus surgical resections compared to autopsy (25) |

MTLE + HS = mesial temporal lobe epilepsy with hippocampal sclerosis

